# *Medicago truncatula Yellow Stripe1-Like3* gene is involved in symbiotic nitrogen fixation

**DOI:** 10.1101/2019.12.23.887448

**Authors:** Rosario Castro-Rodríguez, Isidro Abreu, María Reguera, Lorena Novoa-Aponte, Ana Mijovilovich, Francisco J. Jiménez-Pastor, Javier Abadía, Jiangqi Wen, Kirankumar S. Mysore, Ana Álvarez-Fernández, Hendrik Küpper, Juan Imperial, Manuel González-Guerrero

**Author notes:** Department of Biology. University of Massachusetts. Amherst MA01003. USA. **Corresponding author:** M. González-Guerrero, Centro de Biotecnología y Genómica de Plantas (UPM-INIA), Crta M-40 km39, 28223 Pozuelo de Alarcón (Madrid), Spain.

## Abstract

Symbiotic nitrogen fixation carried out in legume root nodules requires transition metals. These nutrients are delivered by the host plant to the endosymbiotic nitrogen-fixing bacteria living with the nodule cells, a process in which vascular transport is essential. As occurs in root-to-shoot transport, members of the Yellow Stripe-Like (YSL) family of metal transporters should also be required for root-to-nodule metal delivery. The genome of the model legume *Medicago truncatula* encodes for eight YSL proteins, four of them with a high degree of similarity to *Arabidopsis thaliana* YSLs involved in long-distance metal trafficking. Among them, MtYSL3 is a plasma membrane protein expressed by vascular cells in roots and nodules, and by cortical nodule cells. Reducing expression levels of this gene had no major effect on plant physiology when assimilable nitrogen was provided in the nutrient solution. However, nodule functioning was severely impaired, with a significant reduction of nitrogen fixation capabilities. Further, iron and zinc accumulation and distribution changed. Iron was retained in the apical region of the nodule, while zinc became strongly accumulated in the nodule veins in the *ysl3* mutant. These data suggest a role of MtYSL3 in vascular delivery of iron and zinc to symbiotic nitrogen fixation.

**Highlight:** *Medicago truncatula* YSL3 transporter is required for optimal nitrogen fixation in root nodules, mediating iron and zinc distribution in these organs.

## Introduction

Iron, copper, and other transition metals are required at relatively high levels for biological nitrogen fixation, the conversion of N_2_ into NH_4_^+^ carried out by diazotrophic microorganisms (González-Guerrero *et al*., 2014; González-Guerrero *et al*., 2016). These elements act as cofactors of key enzymes mediating the process, such as nitrogenases that directly catalyse the reaction, cytochrome oxidases that provide energy to the reaction and control O_2_ levels, or many of the free radical detoxification enzymes (Appleby, 1984; Rubio *et al*., 2004; Rubio and Ludden, 2005). Therefore, ensuring proper transition metal uptake is critical for any diazotrophic organism. Free-living nitrogen-fixing bacteria use a large battery of siderophores, transition metal transporters, and storage proteins to directly acquire them from the environment (Jurkevitch *et al*., 1992; Yeoman *et al*., 2000; Navarro-Rodríguez *et al*., 2019). In contrast, symbiotic diazotrophs must obtain the required metal nutrients through their host (González-Guerrero *et al*., 2016).

The paradigmatic example of a symbiotic diazotroph are the bacteria known as rhizobia. These organisms colonize the cells of legume root nodules, organs developed to provide the conditions for nitrogen fixation to occur (Downie, 2014). Within the nodule cells, rhizobia are surrounded by plant-derived membranes, the symbiosome membranes, and differentiate into the nitrogen-fixing form, the bacteroids (Kereszt *et al*., 2011). Across the symbiosome membrane, bacteroids deliver the fixed nitrogen while receive photosynthates, phosphate, sulfur, as well as the essential transition elements needed for nitrogen fixation (Udvardi and Poole, 2013). Transition metal nutrients are delivered from the plant root to the nodule through the vasculature and released in the apoplast of the area of bacteroid differentiation (Rodríguez-Haas *et al*., 2013), in a process that resembles metal delivery to shoots (Conte and Walker, 2011). There, different metal transporters introduce these nutrients into the nodule cell cytosol and transfer them across the symbiosome membranes. Several of them have been identified, particularly those proteins located at the host cell plasma membrane and at the symbiosome membrane (Tejada-Jiménez *et al*., 2015; Abreu *et al*., 2017; Tejada-Jiménez *et al.*, 2017; Senovilla *et al*., 2018; Escudero *et al*., 2019a). However, it largely remains to be determined how vascular transport occurs and which proteins are mediating it.

Transition metal loading in the root vasculature is mediated by transporters of the ferroportin and P_1b_-ATPase families (Hussain *et al*., 2004; Andrés-Colás *et al*., 2006; Morrissey *et al*., 2009). Once in the saps, metals are chelated by a collection of soluble molecules, with a prominent role of citrate and nicotianamine (NA). These molecules facilitate metal solubility and prevent oxidative damage (Flis et al., 2016). When metals reach the shoots, they are recovered from the sap as metal-NA complexes and introduced into the cells by members of the Yellow Stripe-Like (YSL) family (Curie *et al.*, 2008). These are a family of plant-specific proteins participating in remobilization of intracellular metal reserves, mediating long-distance metal trafficking and signalling, and in metal uptake from soil by grasses (Curie *et al*., 2001; Waters *et al*., 2006; Conte *et al*., 2013; Kumar *et al*., 2017). In *Arabidopsis thaliana*, AtYSL1 and AtYSL3 are responsible for iron delivery to shoots as well as for signalling iron sufficiency (Waters *et al*., 2006; Kumar *et al*., 2017). Considering the high metal demand of nitrogen fixation (O’Hara, 2001), a large portion of these nutrients has to be delivered to nodules, where similar mechanisms to those reported in shoots would likely be in place. Therefore, it should be expected that metal-NA transporting YSLs similar to AtYSL1 or AtYSL3 are functional in the nodule vasculature. Recent identification of nodule nicotianamine synthases and evidence on their importance for iron homeostasis in nodules supports this hypothesis (Avenhaus *et al*., 2016; Escudero *et al*., 2019b).

Here we report the role of *Medtr3g092090*, MtYSL3, a *Medicago truncatula* orthologue of AtYSL3, highly expressed in nodules and with a vascular localization. Mutation of *MtYSL3* results in a reduction of nitrogenase activity that affects plant growth, the likely consequence of reduced iron and zinc content in nodules and its altered distribution. The data is consistent with a role of MtYSL3 in iron and zinc delivery to nodules.

## Materials and Methods

### Biological Materials and plant growth conditions

*Medicago truncatula* seeds were scarified and surface sterilized following the same protocol described in Tejada-Jiménez *et al*. (2015). After a previous pre-germination step in water agar 0.8% plates during 48h at 22°C, seedlings were planted in sterilized perlite pots and inoculated with *Sinorhizobium meliloti* 2011 or *S. meliloti* 2011 transformed with pHC60 (GFP expressing vector) (Cheng and Walker, 1998) for nodulation assays. Nodules were collected 28 dpi. For non-symbiotic experiments, plants were watered every 2 weeks with solutions supplemented with 2 mM NH_4_NO_3_. In all cases, plants were watered every two days alternating Jenner’s solution with water (Brito *et al*., 1994) and they were grown in a greenhouse under 16 h light / 8 h dark at 25 ºC / 20 ºC conditions. For hairy-roots transformations, *M. truncatula* seedlings were transformed with *Agrobacterium rhizogenes* ARqua1 carrying the appropriate binary vector, as previously described (Boisson-Dernier *et al*., 2001). Transient expression in *Nicotiana benthamiana* were performed by transforming leaves with the plasmid constructs in *Agrobacterium tumefaciens* C58C1 (Deblaere *et al*., 1985). These plants were grown in the greenhouse under the same conditions as *M. truncatula*.

### RNA isolation and quantitative real-time PCR

RNA was extracted from shoos, roots and nodules using Tri-Reagent (Life Technologies), DNase treated and cleaned with RNeasy Minikit (Qiagen, Valencia, CA). One microgram of DNA-free RNA was used to synthesize cDNA by using PrimeScript^TM^ RT Reagent Kit (TAKARA, Kusatsu, Shiga, Japan).

Gene expression was studied by quantitative real-time PCR (StepOne plus, Applied Biosystems) using the Power SyBR Green master mix (Applied Biosystems). Primers used are listed in Table S1. RNA levels were normalized by using the *ubiquitin carboxy-terminal hydrolase* gene as internal standard for *M. truncatula* genes (Kakar *et al.*, 2008).

### β-glucuronidase (GUS) assay

A transcriptional fusion between *MtYSL3* promoter and the β-glucoronidase gene was constructed by amplifying two kilobases upstream of *MtYSL3* start codon using primers indicated on Table S1. This amplicon was inserted into pDONR207 and transferred to pGWB3 (Nakagawa *et al*., 2007) using Gateway technology® (Invitrogen). *M. truncatula* R108 roots were transformed as indicated above. Transformed plants were transferred to sterilized perlite pots and inoculated with *S. meliloti* 2011. GUS activity was determined in 28 dpi plants as described (Vernoud *et al*., 1999).

### Inmunolocalization of MtYSL3-HA

The genomic full sequence of *MtYSL3* including two kilobases upstream of its start codon was amplified using the primers indicated in Suppl. Table 1 and fused with three C-terminal HA epitopes in frame by cloning into the pGWB13 (Nakagawa *et al.*, 2007). Hairy-root transformation was performed as described previously by Boisson-Dernier *et al.* (2001). Transformed plants were inoculated with *S. meliloti* 2011 containing the pHC60 plasmid that constitutively expresses GFP. After 28 dpi, nodules and roots were collected. Fixation and immunohistochemistry protocols were carried out as indicated by Tejada-Jiménez *et al.* (2015).

### Transient expression in *N. benthamiana* leaves

*MtYSL3* coding sequence was cloned into pGWB6 (Nakagawa *et al*., 2007) using Gateway Technology (Invitrogen) resulting in a N-terminal fusion with GFP. This construct, and the plasma membrane marker pBIN *AtPIP2*-CFP (Nelson *et al.*, 2007) were introduced into *A. tumefaciens* C58C1 (Deblaere *et al*., 1985). Transformants were grown in a liquid medium to late exponential phase, centrifuged and resuspended to an OD_600_ of 1.0 in 10 mM MES pH 5.6, containing 10 mM MgCl_2_ and 150 μM acetosyringone. These cells were mixed with an equal volume of *A. tumefaciens* C58C1 expressing the silencing suppressor p19 of Tomato bushy stunt virus (pCH32 35S:p19) (Wood *et al*., 2009). Bacterial suspensions were incubated for 3 h at room temperature and then injected into young leaves of 4 weeks-old *N. benthamiana* plants. Expression of the appropriate construct was analysed after 3 days by confocal laser-scanning microscopy (Leica SP8) with excitation lights of 405 nm for CFP and 488 nm for GFP.

### Nitrogenase activity

The acetylene reduction assay was used to measure the nitrogenase activity (Hardy *et al*., 1968). Wild-type and mutant roots at 28 dpi were introduced separately in 30 ml tubes fitted with rubber stoppers. Each tube contained from three to five roots. Three milliliters of air in each tube was replaced by the same volume of acetylene and subsequently, they were incubated for 30 min at room temperature. Gas samples (0.5 ml) were analyzed in a Shimadzu GC-8A gas chromatograph fitted with a Porapak N column. The amount of ethylene produced was determined by measuring the height of the ethylene peak relative to background. Each point consists of two tubes each measured in triplicate. After measurements, nodules were recovered from roots to measure their weight.

### Metal content measurements

Iron, copper and zinc content were determined in shoots, roots, and nodules 28 dpi. Plant tissues were weighted and mineralized in 15.6 M HNO_3_ (trace metal grade) for 1 h at 80 ºC and overnight at 20 ºC. Digestions were completed with 2 M H_2_O_2_. Samples were diluted in 300 mM HNO_3_ prior to measurements. Element analyses were performed with Atomic Absorption Spectroscopy (AAS) in an AAnalyst 800 (Perkin Elmer), equipped with a graphite furnace. All samples were measured in duplicate.

### Metal localization by micro-X-ray fluorescence (μ-XRF)

A customised benchtop µXRF beamline “M4 Tornado” (Bruker Nano GmbH, Germany) as described in detail by Mijovilovich *et al*. (2020), was used for analysing tissue-level metal distribution in the root nodules. In brief, in this machine the nodules were kept alive in a custom-designed measuring chamber where throughout the measurement they were kept in Jenner’s solution. The measurement was done by excitation with a Rh tube with fibre optic focusing of the beam to 15 µm and filtering of the excitation spectrum with an AlTi filter. A step size of 8 µm was applied to yield a 2x oversampling. The measured µXRF spectra in each pixel of the µXRF maps were deconvoluted using the software supplied with the Tornado. The net counts in the resulting element distribution maps were recalculated to mM concentrations according to a certified liquid standard (standard solution VI, Merck KGaA Darmstadt Germany) in a cuvette of the thickness of an average nodule. Colour scales were assigned to the quantified data using ImageJ, after which they were converted from 16bit to RGB format for assembling the figure using PhotoImpact X3 (Corel Corporation, Ottawa, Canada).

### NA content determination

Nicotianamine was extracted as previously described (Banakar *et al.*, 2017) with some modifications. Briefly, nicotianamine was extracted from approximately 50 mg of nodules, and frozen grinded in 400 µl miliQ water spiked with nicotyl-lysine (150 µM final concentration) as internal standard. Samples were homogenized in a mixer mill (Retsch MM300, Retsch) during 5 min at 30 sec^−1^ frequency, and then centrifuged at 12000 g for 10 minutes at 4 ºC. Supernatant was then passed through a 3 kDa cutoff centrifugal filter (cellulose Amicon®, Merck) (1 h at 14000 g at 4 ºC) and dried under vacuum (1.5 h at 40 °C). Dry residues from shoots were dissolved in 20 µl of miliQ water, whereas root-and-nodule dry residues were dissolved in 15 µl. Then 5 µl aliquots were mixed with EDTA (final concentration 8.33 mM) to dissociate potential nicotianamine-metal complexes, and 50 % (v/v) mobile phase A (see below) to favor chromatographic separation. The mixture was filtered through 0.45 polyvinylidene fluoride (PVDF) ultrafree-MC centrifugal filter devices (Merck) before analysis.

Nicotianamine levels were determined by high-performance liquid chromatography electrospray ionization time-of-flight mass spectrometry (HPLC-ESI-TOF-MS) as described by Banakar et al. (2017). The samples were fractionated using an Alliance 2795 HPLC system (Waters) and µLC column (SeQuant ZIC®-HILIC, 15 cm x 1 mm internal diameter, 5 µm, 200 Å, Merck), with a mobile phase consisting of solvent A (10 % 10 mM ammonium acetate pH 7.3 90 % acetonitrile) and solvent B (80 % 30 mM ammonium acetate pH 7.3 20 % acetonitrile) at a flow rate of 0.15 ml min^−1^. The gradient program started at 100 % (v/v) solvent A for 3 min, and then decreased linearly to 30 % (v/v) solvent A over the next 7 min, then remained for 7 min at 30 % (v/v) solvent A, and then returned to the initial conditions over the next 8 min. The column was then allowed to stabilize for 10 min at the initial conditions before proceeding to the next injection. The total HPLC run time was 35 min, the injection volume was 10 µl and the auto sampler and column temperatures were 6 ºC and 30 ºC, respectively. The HPLC was coupled to the MicrOTOF mass spectrometer (Bruker Daltonics) equipped with an ESI source. The operating conditions were optimized by the direct injection of 100 µM solutions of nicotianamine standard at a flow rate of 180 µl h^−1^. Mass spectra were acquired in negative ion mode over the 150–700 mass-to-charge (*m/z*) ratio range. The mass axis was calibrated externally using Li–formate adducts (10 mM LiOH, 0.2 % (v/v) formic acid and 50% (v/v) 2-propanol). Bruker Daltonik software packages micrOTOF Control v2.2, HyStar v3.2 and Data Analysis v4.0 were used to control the MS, HPLC interface and for data processing, respectively. Nicotianamine (Toronto Research Chemicals) calibration curve were prepared with nycotyl-lysine as internal standard.

### Bioinformatics

To identify *M. truncatula* YSL family members, BLASTN and BLASTX searches were carried out in the *M. truncatula* Genome Project site (http://www.jcvi.org/medicago/index.php). Protein sequences for tree construction were obtained from Phytozome (https://phytozome.jgi.doe.gov/pz/portal.html), Uniprot (http://www.uniprot.org/blast/) and from NCBI (https://blast.ncbi.nlm.nih.gov/Blast.cgi?PAGE=Proteins): *Medicago truncatula* MtYSL1 (Medtr1g077840); MtYSL2 (Medtr1g007540); MtYSL3 (Medtr3g092090); MtYSL4 (Medtr1g007580); *Arabidopsis thaliana* AtYSL1 (At4g24120), AtYSL2 (At5g24380), AtYSL3 (At5g53550), *Oryza sativa* OsYSL15 (Os02g0650300), OsYSL16 (Os04g0542800); *Zea mays* ZmYS1 (Zm00001d017429), ZmYSL2 (Zm00001d025977), *Brachypodium distachyon* BdYS1A (BRADI_3g50267), BdYS1B (BRADI_3g50263), BdYSL2 (BRADI_3g50260), BdYSL3 (BRADI_5g17230).

Trees were constructed from a ClustalW multiple alignment of the sequences (http://www.ebi.ac.uk/Tools/msa/clustalw2), then analyzed by MEGA7 (Kumar *et al.*, 2016) using a Neighbour-Joining algorithm with bootstrapping (1,000 iterations). Unrooted trees were visualized with FigTree (http://tree.bio.ed.ac.uk/software/figtree).

The topology modelling were performed using the visualization software PROTTER (http://wlab.ethz.ch/protter/start/), which includes the transmembrane region prediction software Phobius.

### Statistical tests

Data were analyzed with Student’s unpaired t test to calculate statistical significance of observed differences. Test results with p-values lower than 0.05 were considered as statistically significant.

## Results

### *MtYSL3* is highly expressed in the nodule vasculature

The proteins of the YSL family cluster in four groups, with group I being the best characterized (Yordem *et al*., 2011). It includes AtYSL1, 2 and 3, founding protein *Zea mays* YS1, and four *M. truncatula* YSLs (MtYSL1-4, *Medtr1g077840, Medtr1g007540, Medtr3g092090, Medtr1g007580*, respectively) (Fig. 1A). Among the *M. truncatula* group I YSLs, expression of *MtYSL4* was not detected in any of the organs tested (Fig. S1), while the other three were expressed in shoots and roots from inoculated and non-inoculated plants, as well as in nodules. (Fig. 1B, Fig. S1). *MtYSL1* transcripts were more abundant in shoots than anywhere else in the plants. *MtYSL2* and *MtYSL3* transcription was more intense in nodules, being the latter the most highly expressed, approximately four times higher in nodules than in any other plant organ. The inoculation with *S. meliloti* did not result in significant transcriptional changes in shoots or roots compared to non-inoculated nitrogen-fertilized plants.

**Fig. 1.**
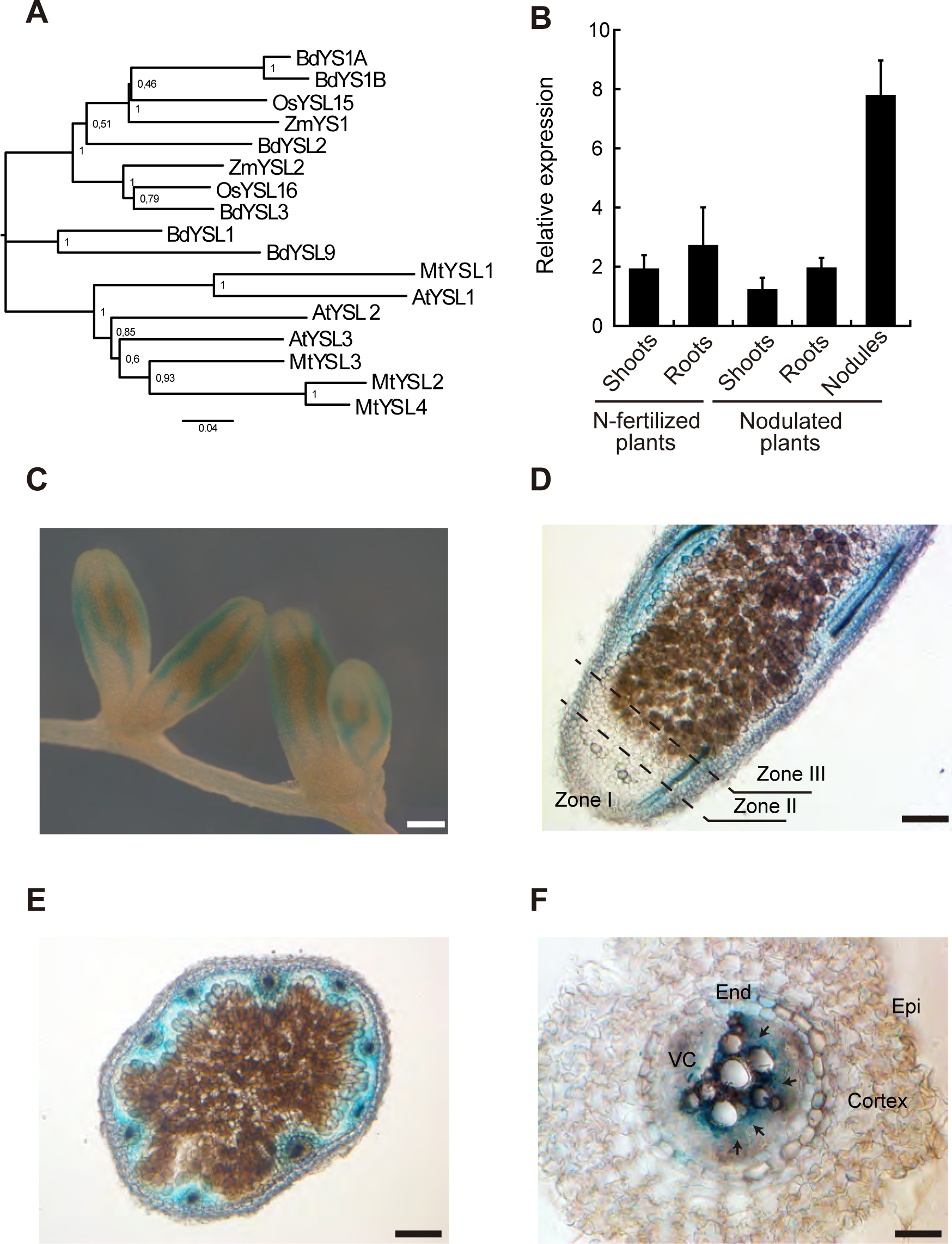
*MtYSL3* is highly expressed in nodule and root vasculature. **A**) Phylogenetic tree of the Group I YSLs transporters, MtYSL1-4 (*Medtr1g077840, Medtr1g007540, Medtr3g092090* and *Medtr1g007580* respectively) and their representative homologues in *Zea mays*, *Oryza sativa*, *Brachypodium distachyon* and *Arabidopsis thaliana*. **B**) Gene expression relative to the internal standard gene *ubiquitin carboxyl-terminal hydrolase* in shoots, roots and nodules of nitrogen-fertilized plants and nodulated plants. Data are the mean ± SE of five independent experiments. **C**) Histochemical staining of GUS activity in 28 dpi root and nodules of *M. truncatula* plants transformed with containing the *MtYSL3-promoter::gus*. Scale bar = 500 μm. **D**) Longitudinal section of a GUS-stained 28 dpi nodule expressing *gus* under *MtYSL3* promoter. Scale bar = 200 μm. **E**) Cross section of a GUS-stained nodule expressing *gus* under the *MtYSL3* promoter. Scale bar = 200 μm. **F**) Cross section of a GUS-stained root expressing *gus* under the *MtYSL3* promoter. End: endodermis, Epi: epidermis, VC: vascular cylinder. Scale bar = 50 μm.

To locate the tissue expression of *MtYSL3*, the 2 kb region upstream of *MtYSL3* was used to drive the expression of the β-glucuronidase (*gus*) gene. After 28 days post-inoculation (dpi), roots and nodules were incubated with X-gluc to visualize the GUS activity. The staining pattern was consistent with a vascular expression of *MtYSL3* in both organs (Fig. 1C). Longitudinal sections of those nodules revealed a peripheral distribution of the signal, associated to the vasculature, and no expression in the inner nodule region (Fig. 1D). This was also supported by nodule cross-section images (Fig. 1E). In addition, some GUS activity was observed in these sections in cortical nodule cells, although at much lower intensity than in the vasculature. In roots, *MtYSL3* expression was confined to the endodermis and inner vascular layers (Fig. 1F).

Immunolocalization of epitope-tagged MtYSL3 supports the GUS activity assays. Three hemagglutinin (HA) tags were fused to the C-terminus of the protein and expressed under its own promoter region. MtYSL3-HA localization was visualized using a primary anti-HA mouse antibody and an Alexa594-conjugated anti-mouse antibody. The transformed plants were inoculated with a strain of *S. meliloti* that constitutively expresses GFP. The HA epitope of MtYSL3-HA was detected in the vasculature of the nodule and in cortical cells (Fig. 2A and B). Closer detail of the vascular region showed colocalization with the autofluorescence pattern of the Casparian strip, indicating that MtYSL3-HA was located in the endodermis (Fig. 2C). In roots, MtYSL3-HA was observed in the endodermis and in inner vascular cells, very likely the xylem parenchyma (Fig. 2D). The peripheral distribution of the Alexa594 signal was indicative of a plasma membrane distribution. To test this possibility, *N. benthamiana* leaves were co-agroinfiltrated with a plasmid constitutive expressing *MtYSL3* fused to GFP and plasma membrane marker AtPIP2 fused to CFP. As shown in Fig. 2E, both the GFP and the CFP signal colocalized. Controls did not show any autofluorescence in the Alexa594 channel in the conditions tested (Fig. S2).

**Fig. 2.**
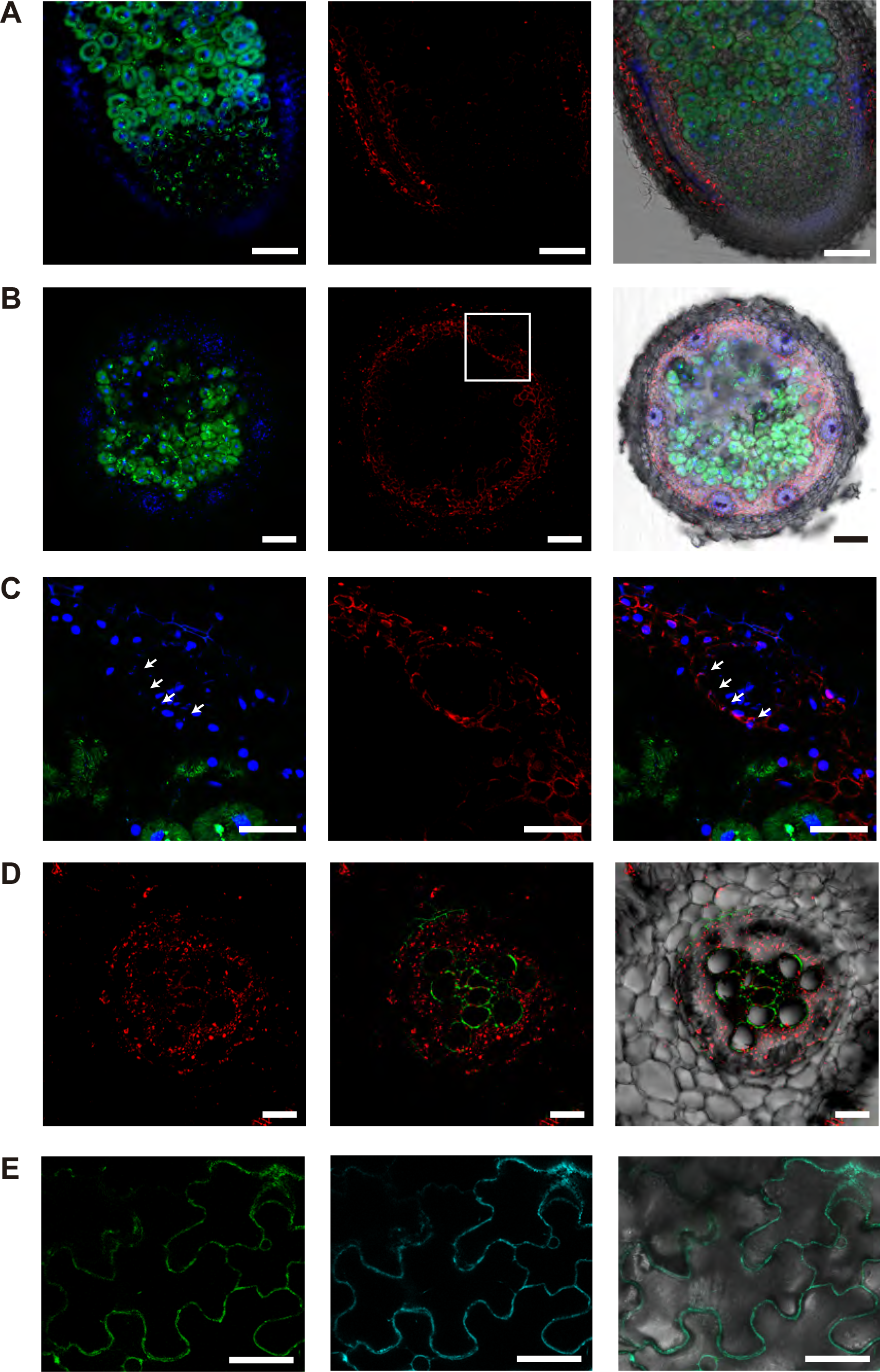
MtYSL3-HA is located in the plasma membrane of endodermal cells in roots and nodules, in root xylem parenchyma and in the nodule cortical cells. **A**) Longitudinal and **B**) cross-section of 28 dpi *M. truncatula* nodules colonized with *S. meliloti* constitutively expressing GFP (green) and transformed with a vector expressing the fusion MtYSL3-HA under the regulation of its endogenous promoter. MtYSL3-HA localization was determined using an Alexa 594-conjugated antibody (red). DNA was stained with DAPI (blue). Left panels show the overlay of GFP and DAPI channels; centre panels, the Alexa594 channel; and right panels he overlay of GFP, Alexa594 and DAPI channels with the brightfield image. Scale bars = 200 μm (A) or 50 μm (B). **C**) Magnification of the vascular bundle within the boxed region indicated in (B). Left panels show the overlay of GFP and DAPI channels; centre panels, the Alexa594 channel; and right panels he overlay of the three channels. Arrows indicate the position of the autofluorescence signal of the Casparian strip. Scale bars = 50 μm. **D**) Cross section of a *M. truncatula* root transformed with a vector expressing the fusion MtYSL3-HA under regulation of its endogenous promoter located with an Alexa594-conjugated antibody (red, left panel). Centre panel shows it colocalization with the autofluorescence signal of lignin (green). Right panel shows the overlaid images with the transillumination channel. Scale bars = 50 μm. **E**) Transient expression of MtYSL3-GFP and AtPIP2-CFP in *N. benthamiana* leaves. Left panel shows the localization of MtYSL3 fused to GFP (green) in tobacco cells. Middle panel shows the localization of plasma membrane marker AtPIP2 fused to CFP in the same cells. Right panel is the overlay of the two previous channels together with the bright field image. Scale bars = 50μm.

### MtYSL3 is involved in symbiotic nitrogen fixation

To determine the role of MtYSL3 in *M. truncatula* physiology, two *Tnt1* insertion lines were obtained from the Noble Research Institute (Tadege *et al.*, 2008). NF17945 (*ysl3-1*) presents an insertion in position +342, within the first exon of the gene (Fig. 3A). NF12068 (*ysl3-2*) is inserted in the promoter region of *MtYSL3*, in position −19. While in both cases *MtYSL3* expression was detected, these *Tnt1* lines showed a severe reduction of *MtYSL3* transcript compared to wild type plants (Fig. 3B). Transposon insertion in *ysl3-1* resulted in an altered splicing that left a 30 nucleotide insertion of the *Tnt1* sequence in *MtYSL3* mRNA. As result, five amino acids were mutated (Y115D; S116D; I117V; A118H; G120L) and four more added between amino acid 118 and 119 (LIEE). These changes occurred in a predicted transmembrane domain, and would likely disrupt this region, as indicated by the transmembrane region prediction software Phobius (http://phobius.sbc.su.se/) (Fig. S3). Loss of transmembrane domains would cause a major disruption on the functionality of any membrane protein, and thus *ysl3-1* has been considered as a loss-of-function mutant, while *ysl3-2* would be a knock-down line.

**Fig. 3.**
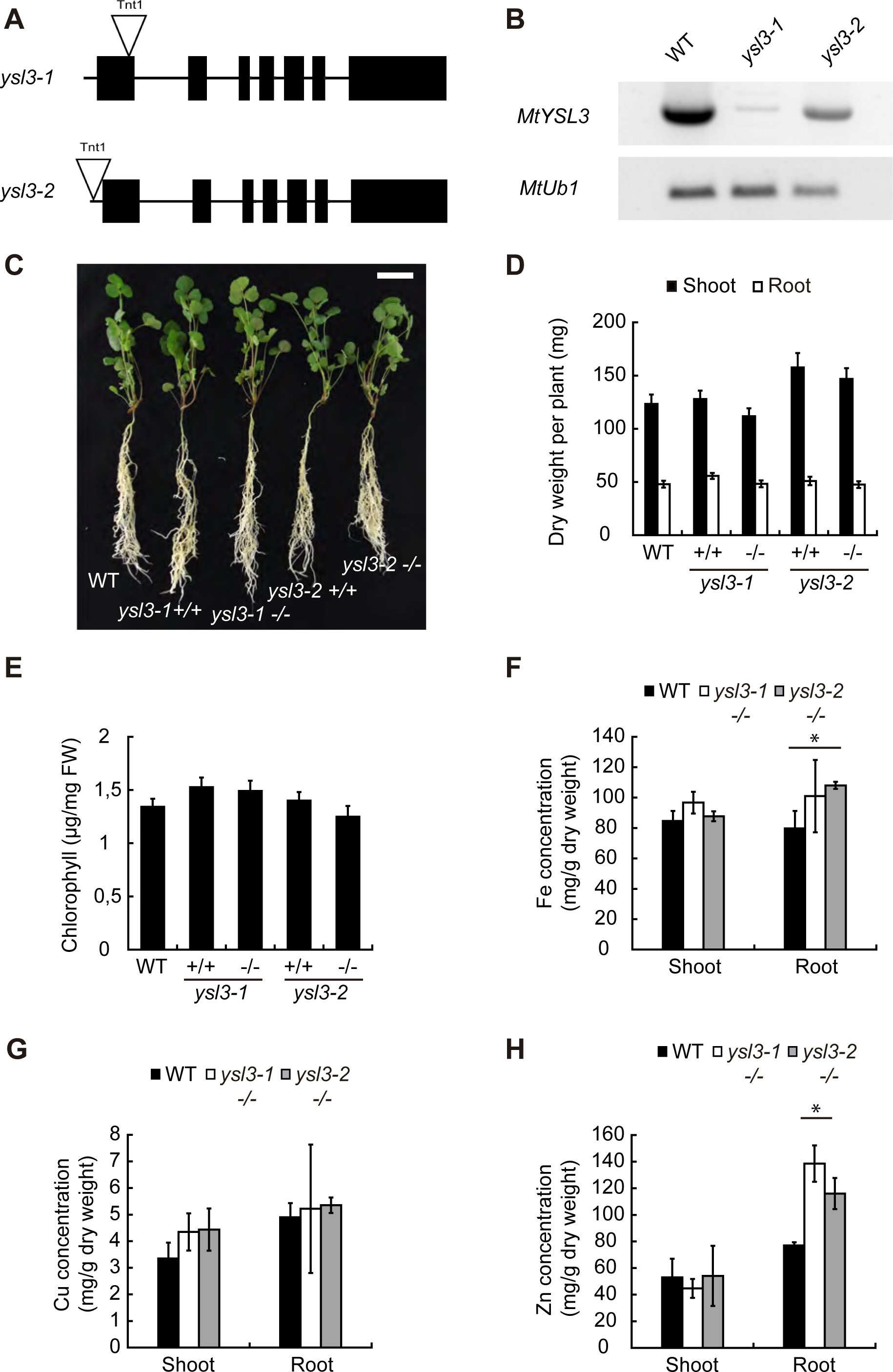
*MtYSL3* does not play an important role under non-symbiotic conditions. **A**) Position of the *Tnt1* insertion site for *ysl3-1* (NF17945) and for *ysl3-2* (NF12068). **B**) RT-PCR of *MtYSL3* expression in 28 dpi nodules in wild-type (WT), *ysl3-1* and *ysl3-2 M. truncatula* lines. Expression of *ubiquitin carboxyl-terminal hydrolase* was used as positive control. **C**) Growth of representative WT, *ysl3-1* and *ysl3-2* plants. +/+ indicates *ysl3* segregants with two wild-type copies of *MtYSL3*, while −/− indicate that both copies have the *Tnt1* insertion. Bar = 3 cm. **D**) Dry weight of shoots and roots of 28 dpi plants. Data are the mean ± SE (n = 20-60 plants). **E**) Chlorophyll content of WT, *ysl3-1 +/+, ysl3-1 −/−, ysl3-2 +/+* and *ysl3-2 −/−.* Data are the mean ± SE of ten sets of four-five pooled plants. **F**) Iron content in roots and shoots of WT, *ysl3-1* −/− and *ysl3-2* −/− plants. Data are the mean ± SE of three pools of four-five plants. **G**) Copper content in roots and shoots of WT, *ysl3-1* −/− and *ysl3-2* −/− plants. Data are the mean ± SE of three pools of four-five plants. **H**) Zinc content in roots and shoots of WT, *ysl3-1* −/− and *ysl3-2* −/− plants. Data are the mean ± SE of three pools of four-five plants.

Under non-symbiotic conditions, when the plants were not inoculated with rhizobia but fertilized with ammonium nitrate, no significant differences were observed in plant growth and biomass production between wild type plants, *Tnt1* segregants with two wild type copies of *MtYSL3* (+/+ lines), or segregants with both *MtYSL3* copies mutated (-/-lines) (Fig. 3C and D). No significant differences were observed in either total chlorophyll content (Fig. 3E) or iron concentration in shoots either (Fig. 3F). However, *ysl3* plants had a trend to accumulate more iron in roots, significatively so in the *ysl3-2* allele. Copper levels were not significantly different in shoots or roots (Fig. 3G), while zinc concentrations were significantly higher in the roots of both *MtYSL3* mutants (Fig. 3H). Unlike *A. thaliana* orthologues (Waters *et al*., 2006), *MtYSL3* mutant plants did not show any significant reduction in fertility, as indicated by the number of pods per plant and of seeds within them (Fig. S4).

However, when the nitrogen was provided by the endosymbiotic rhizobia within their root nodules, *ysl3-1* and *ysl3-2* −/− lines had a smaller growth that their wild type segregants (Fig. 4A). This was also shown when comparing the dry weight of loss-of-function *ysl3-1* −/− to the wild type segregant, with significantly lower biomass production (Fig. 4B). While there were no significant changes in the number of nodules per plant (Fig. 4C), both mutant lines had approximately 60% of the nitrogenase activity of the wild type control or +/+ segregant lines (Fig. 4D). This could be due to the reduced iron content in nodules of *ysl3-1* and *ysl3-2* plants (Fig. 4E). While copper concentration did not significantly change in these organs, *ysl3-2* nodules had less zinc (Fig. 4F and G). However, copper was more concentrated in *ysl3-1* shoots. Similarly to non-inoculated plants, there was no significant difference in pod or seed production in these plants compared to their control (Fig. S5).

**Fig. 4.**
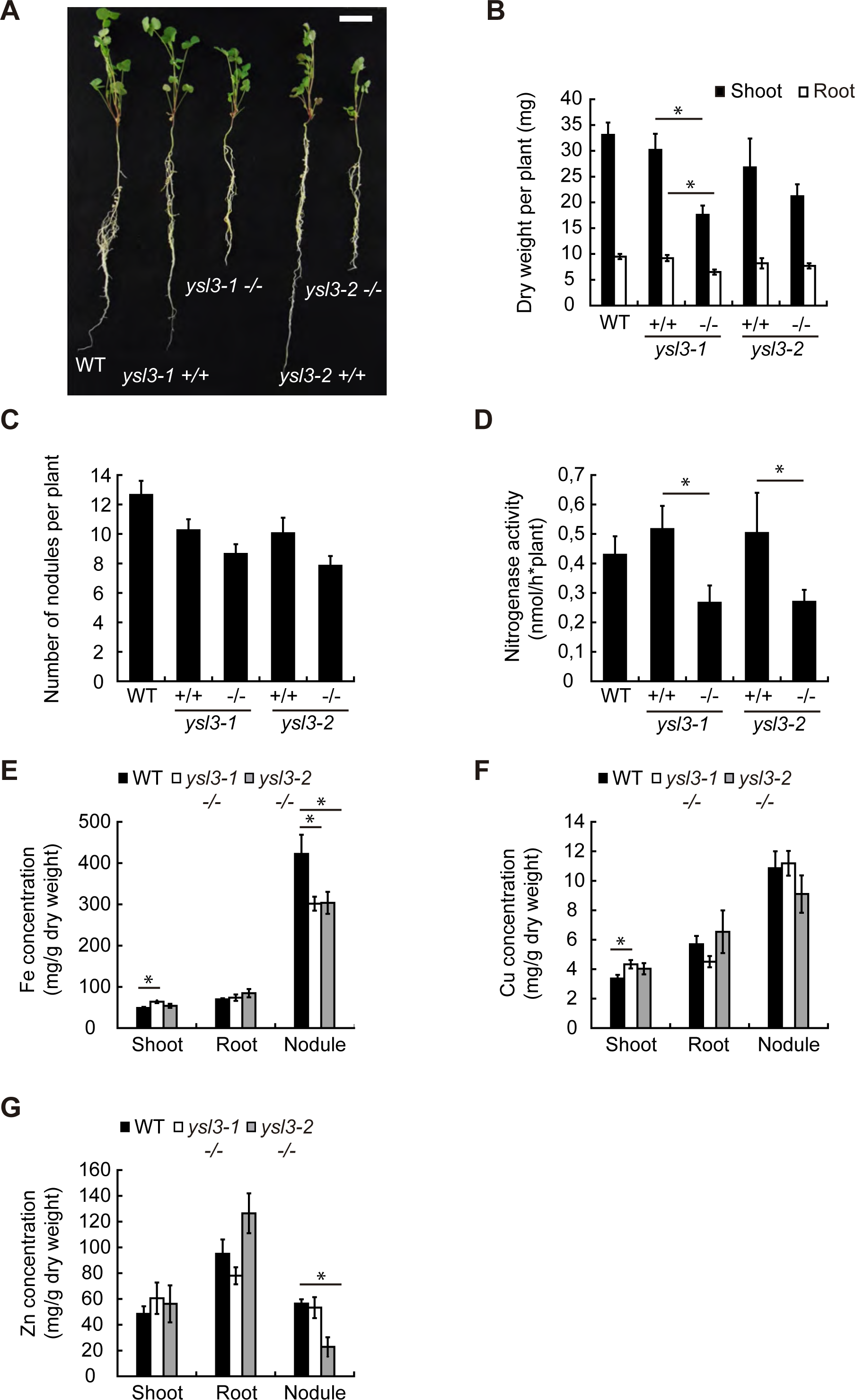
*MtYSL3* participates in symbiotic nitrogen fixation. **A**) Representative WT, *ysl3-1* and *ysl3-2* plants. +/+ indicates *ysl3* segregants with two wild-type copies of *MtYSL3*, while −/− indicate that both copies have the *Tnt1* insertion. Bar = 3 cm. **B)** Dry weight of shoots and roots of 28 dpi plants. Data are the mean ± SE (n = 10-40 plants). **C)** Number of nodules per plant in WT and mutant lines. Data are the ± SE (n = 10-40 plants). **D)** Nitrogenase activity in 28 dpi nodules from WT, *ysl3-1*, and *ysl3-2* plants. Data are the mean ± SE (n = 5-15 sets of pooled plants). **E)** Iron content in shoots, roots and nodules of WT, *ysl3-1* −/− and *ysl3-2* −/− plants. Data are the mean ± SE of three pools of four-five plants. **F)** Copper content in shoots, roots and nodules of WT, *ysl3-1* −/− and *ysl3-2* −/− plants. Data are the mean ± SE of three pools of four-five plants. **G**) Zinc content in roots and shoots of WT, *ysl3-1* −/− and *ysl3-2* −/− plants. Data are the mean ± SE of three pools of four-five plants.

### MtYSL3 silencing affects iron and zinc distribution

The reduced iron content in *ysl3* nodules and lower nitrogenase activity could be the result of less iron being delivered to the fixation zone. To test this possibility, we carried out X-ray fluorescence tests in nodules from wild type and *ysl3-1* nodules (Fig. 5A). While a typical wild type nodule has less iron in the apical region relative to the fixation zone (Fig. 5B), *ysl3-1* nodules had the opposite, indicating that not enough iron would be reaching the fixation zone. Furthermore, while significant changes in nodule zinc concentration was observed only in *ysl3-2* nodules, zinc distribution was also affected in *ysl3-1* nodules. X-ray fluorescence data showed that this nutrient accumulated at much larger levels in the *ysl3-1* nodule vessels than in wild-type ones (Fig. 5A, C). Although metal-nicotianamine would be the likely substrate of MtYSL3, no significant differences in nicotianamine content in *ysl3-1* nodules were observed compared to the wild type (Fig. S6).

**Fig. 5.**
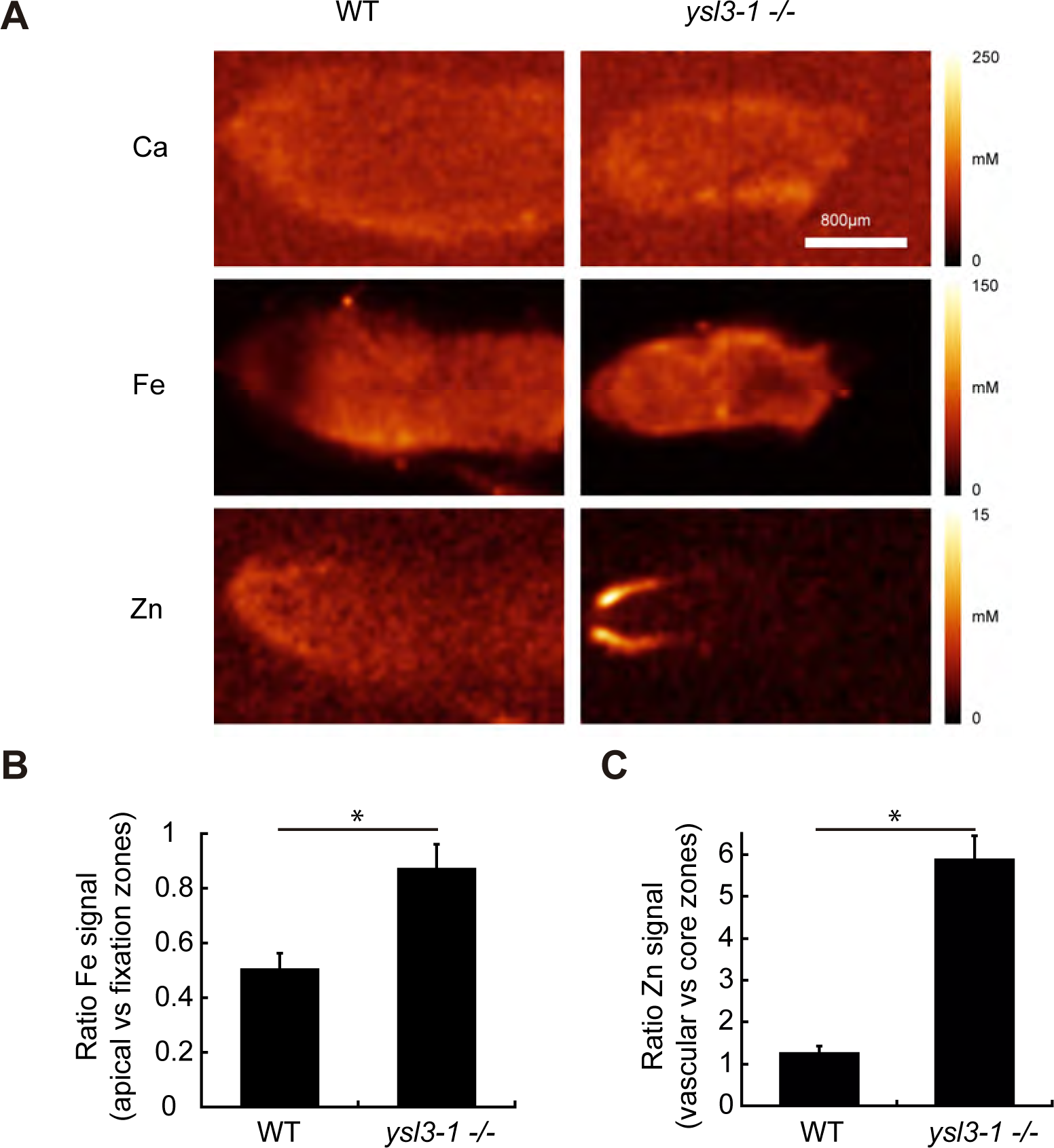
Mutation of *MtYSL3* alters nodule iron and zinc distribution. A) X-ray fluorescence localization of calcium (top panels), iron (centre panels) and zinc (lower panels) in representative 28 dpi nodules from WT and *ysl3-1 −/−* plants. **B**) Ratio of iron concentration in the apical *vs* the fixation zone in 28 dpi nodules from WT and *ysl3-1 −/−* plants. Data are the mean ± SE (n = 4-5 nodules). **C**) Ratio of zinc concentration in the nodule core *vs* the vasculature in 28 dpi nodules from WT and *ysl3-1 −/−* plants. Data are the mean ± SE (n = 4-5 nodules).

## Discussion

Transition metals are essential plant nutrients (Marschner, 2011). Typically, the main plant metal sinks are in the leaves, where these elements participate in the electron transport chains in photosynthesis and oxidative respiration, among several other processes; and in the seeds, where they are critical for embryo development and germination (Kobayashi and Nishizawa, 2012; Yruela, 2013; Ibeas *et al.*, 2017). Consequently, plants dedicate a large effort to ensure metal translocation from roots to leaves and seeds, which includes the participation of Arabidopsis YSL1 and YSL3 (Curie *et al*., 2008; Conte *et al*., 2013). However, this is more challenging in legumes when they are in symbiosis with rhizobia. Due to the large amounts of metalloenzymes participating in symbiotic nitrogen fixation (Brear *et al*., 2013; González-Guerrero *et al.*, 2014), nodules are a major metal sink. A third of the total plant iron content and a quarter of the total copper and zinc are concentrated in nodules, which in the case of *M. truncatula*, represents around 5% of the total plant biomass (Tejada-Jiménez *et al.*, 2015; Abreu *et al*., 2017; Senovilla *et al*., 2018). Therefore, legumes have to direct large quantities of metals not only to shoots, but also to nitrogen-fixing nodules. In this task, it is likely that YSL proteins similar to *A. thaliana* YSL1 and YSL3 would participate.

MtYSL3 is one of the four clade I YSLs in *M. truncatula*. It is closely related to *A. thaliana* YSL1 and YSL3, both being responsible for long-distance metal delivery (Walker and Waters, 2011). As it was the case for the *A. thaliana* orthologues, *MtYSL3* is expressed in the vasculature, both in roots and in nodules. This was confirmed by immunolocalization of a HA-tagged protein using confocal microscopy. Moreover, MtYSL3-HA had a plasma membrane localization in endodermal and in root xylem parenchyma cells. In nodules, some expression could also be detected in nodule cortical cells, those in the exterior of the nodule. This localization would be consistent with a role in vascular transition metal transport, as well as metal uptake by nodule cortical cells.

In non-inoculated plants, MtYSL3 does not seem to play a critical role by itself. There are no major changes in biomass production, leave chlorosis, or plant fertility. There is a significant accumulation in roots of zinc in both mutant alleles and of iron in one of them. But this had no effect on total shoot metal concentrations. This is consistent with the reported functional redundancy of the YSL family. Likely candidates to complement MtYSL3 would be MtYSL1 (orthologue to AtYSL1 and expressed primarily in shoots and roots) or MtYSL2 (similar expression profile as *MtYSL3* but at lower levels). In contrast, when *M. truncatula* is nodulated, mutating or simply silencing *MtYSL3* expression results in a 40% reduction of nitrogenase activity, with a significant reduction on biomass production when MtYSL3 is inactivated. Moreover, iron and zinc accumulation and distribution are affected in the *MtYSL3* mutants. Less iron reaches *ysl3-1* and *ysl3-2* nodules, and it is less abundant in the fixation zone what would result in less iron being available for nitrogenase cofactor synthesis. Zinc is retained in the nodule vessels what would result in lower amounts available for nodule functioning, although the precise role of zinc in nodule functioning is not yet determined.

Therefore, it could be proposed that MtYSL3 would be participating in iron and zinc delivery to nodules, considering that iron levels were reduced in nodules, and zinc became trapped in the veins of *ysl3*. However, MtYSL3 is likely not the only transporter mediating this process. Any major disruption of iron, copper, or zinc delivery to nodules results in a more severe reduction of nitrogenase activity, the presence of white, non-functional nodules, and/or a reduction in their size (Tejada-Jiménez *et al*., 2015; Abreu *et al*., 2017; Tejada-Jiménez *et al*., 2017; Kryvoruchko *et al*., 2018; Senovilla *et al*., 2018). In contrast, the *ysl3* phenotype was milder. This would indicate that another, yet-to-be-determined transporter might be involved in metal delivery from the vasculature. Based on the available transcriptomic data, it is unlikely that a clade I YSL protein would be carrying out this role to a larger degree than MtYSL3, considering their lower expression levels in nodules.

The localization of MtYSL3 also indicates additional roles to delivering metals to nitrogen-fixing nodule cells. If this were its unique function, *MtYSL3* would only be expressed in the vascular region in the infection/differentiation zone of the nodule, since this is the region where plant-delivered transition metals are released (Rodríguez-Haas *et al.*, 2013). In contrast, as shown for the molybdate vascular transporter MtMOT1.2 (Gil-Díez *et al*., 2019), MtYSL3 is located along the whole nodule vessels, including the fixation zone. Moreover, no polar distribution of the transporter was observed, what suggests that mass effect would drive the net direction of the MtYSL3 substrate, as proposed for MtMOT1.2. This would be compatible with MtYSL3 being involved in recovering iron and zinc from the apoplast in Zone III, a role of particular importance when considering the prevalent low metal bioavailability in soils (Kim and Guerinot, 2007; Alloway, 2008). However, this metal recovery capability would not be essential either, since none of the *MtYSL3* mutants had any alteration in fertility, as was reported when both *AtYSL1* and *AtYSL3* were mutated (Waters *et al.*, 2006). This is in contrast to the proposed role of nicotianamine in metal recycling from senescent nodules (Hakoyama *et al.*, 2009). All this evidence also indicates the existence of a redundant protein that would be carrying out this function together with MtYSL3.

In addition to the vasculature, MtYSL3 is also located in the cortical nodule cells, although expressed at lower levels. There, it could be introducing metal-NA complexes (including iron-NA) into these cells. This is in contrast to rhizobia-infected cells, which introduce iron as Fe^2+^ through a Nramp protein (Tejada-Jiménez *et al*., 2015), playing citrate an important role in its solubility in the apoplast (Takanashi *et al*., 2013; Kryvoruchko *et al*., 2018). This would indicate the existence of different iron pools to separate a limiting nutrient with different physiological functions. Previous data on zinc transporter MtZIP6, only expressed in rhizobia-infected cells, also hints to at least a partial tissue specialization of metal transport (Abreu *et al*., 2017). However, work on *in situ* metal speciation analyses should shed some light into this possibility once synchrotron-based X-ray absorption near-edge structure reach the required sensitivity and resolution and open the study of the mechanisms of intertissued metal sorting in nodules.

In summary, our data indicates that MtYSL3 is involved in vascular transition metal delivery, likely iron and zinc, to nitrogen fixing nodule cells, as well as uptake by cortical cells. Our data also suggests that at least an additional metal transporter is performing analogous functions, since the *ysl3* phenotype is relatively mild. Future work will be directed towards unveiling these additional transporters.

## Supplementary Data

**Fig. S1. Group I *M. truncatula* YSLs expression**.

**Fig. S2**. **Autofluorescence control for Alexa594 signal**.

**Fig. S3. Effect of *ysl3-1 Tnt1* insertion in MtYSL3 topology**.

**Fig. S4**. **Effect of *MtYSL3* mutation in plant fertility under non-symbiotic conditions**.

**Fig. S5**. **Effect of *MtYSL3* mutation in plant fertility under symbiotic conditions.**

**Fig. S6. Nicotianamine content in 28 dpi wild type, *ysl3-1*, and *ysl3-2* nodules**.

**Table S1. Primers used in this study.**

## Acknowledgements

This research was funded by a European Research Council Starting Grant (ERC-2013-StG-335284) and a Ministerio de Economía y Competitividad (MINECO) grant (AGL2015-65866-P), to MGG, and a MINECO grant (AGL2016-75226-R) to JA and AA-F. RC-R was supported by a Formación del Personal Investigador fellowship (BES-2013-062674). IA is recipient of a Juan de la Cierva-Formación postdoctoral fellowship from Ministerio de Ciencia, Innovación y Universidades (FJCI-2017-33222). Development of *M. truncatula Tnt1* mutant population was, in part, funded by the National Science Foundation, USA (DBI-0703285) to KSM. AM and HK and the µXRF measurements were supported by the Ministry of Education, Youth and Sports of the Czech Republic with co-financing from the European Union (grant "KOROLID", CZ.02.1.01/0.0/0.0/15_003/0000336) and the Czech Academy of Sciences (RVO: 60077344). We would also like to acknowledge the other members of laboratory 281 at Centro de Biotecnología y Genómica de Plantas (UPM-INIA) for their support and feedback in preparing this manuscript.

