## Supporting information for "*Medicago truncatula Yellow Stripe1-Like3* gene is involved in symbiotic nitrogen fixation"

### FIGURE S1

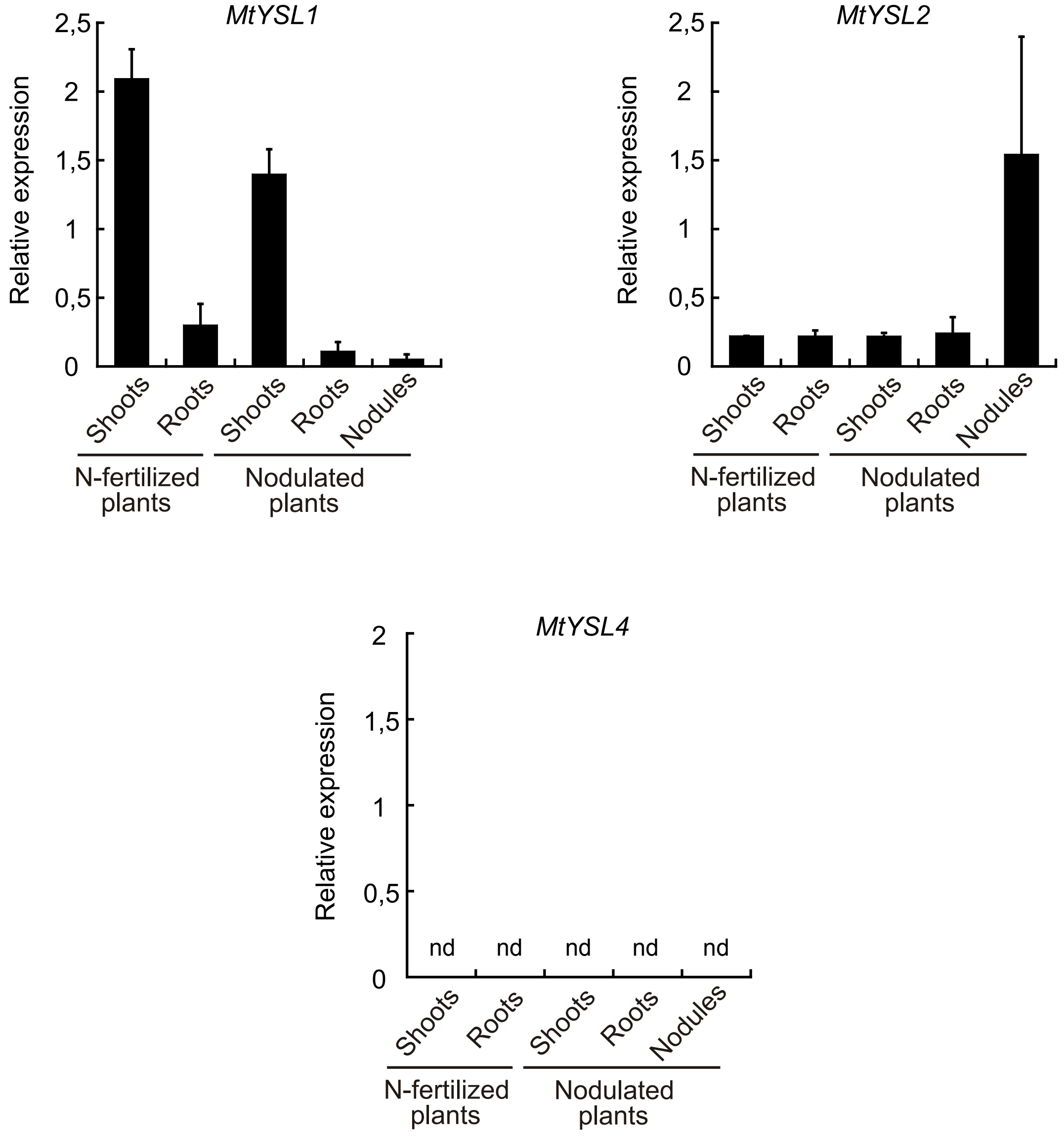


**Fig. S1. Group I *M. truncatula* YSLs expression.** *MtYSL1, 2* and *4* expression relative to internal standard gene *ubiquitin carboxyl-terminal hydrolase*. Data are de mean ± SE of three independent experiments. Nd indicates non-detected expression.

#### FIGURE S2

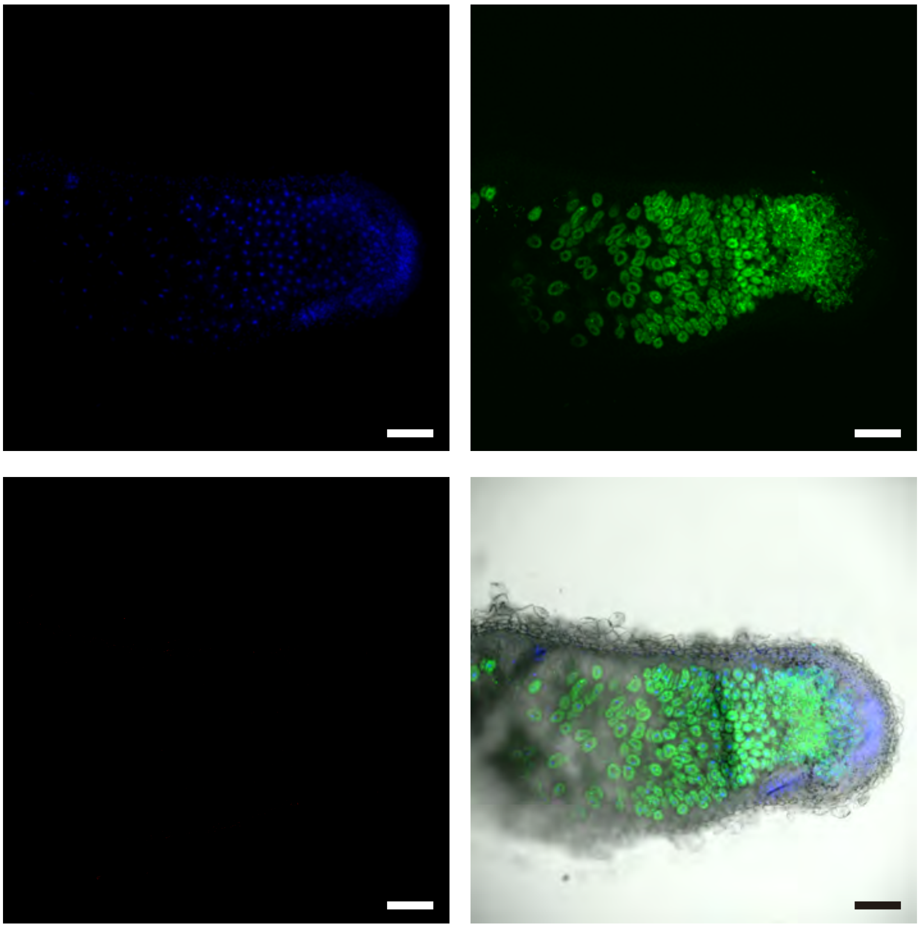

**Fig. S2. Autofluorescence control for Alexa594 signal**. Inmunolocalization of MtYSL3-HA nodule sections from 28 dpi *M. truncatula* plants expressing *MtYSL3-HA*. Sections were treated as indicated for confocal microscopy, but without adding the Alexa594-conjugated antibody. Upper left panel, DAPI-stained DNA (blue); upper right panel, GFP-expressing *S. meliloti*; lower left panel, Alexa594 emission detected in the same conditions as in Figure 2; lower right panel, overlay of the other three panels with the transillumination image obtained. Scale bars = 160 μm.

#### FIGURE S3

**A**

```
MtYSL3 protein   MDTRSNEEEREIENHHIEEGQVAMDEELNRIAPWRKQITVRGLIASLIIGIIYSVIVMKL
ysl3-1 mutant    MDTRSNEEEREIENHHIEEGQVAMDEELNRIAPWRKQITVRGLIASLIIGIIYSVIVMKL
                 ************************************************************

MtYSL3 protein   NLTTGLVPNLNVSAALLGFVFIRSWTKILSKANIVSAPFTRQENTIIQTCAVACYSIA--
ysl3-1 mutant    NLTTGLVPNLNVSAALLGFVFIRSWTKILSKANIVSAPFTRQENTIIQTCAVACDDVHLI
                 ******************************************************  .:

MtYSL3 protein   --VGGGFGSYLLGLNRRTYEQAGIDTPGNTPGSTKEPAIGWMTAFLFVTSFVGLLALVPI
ysl3-1 mutant    EEVLGGFGSYLLGLNRRTYEQAGIDTPGNTPGSTKEPAIGWMTAFLFVTSFVGLLALVPI
                    * ********************************************************
```

**B**

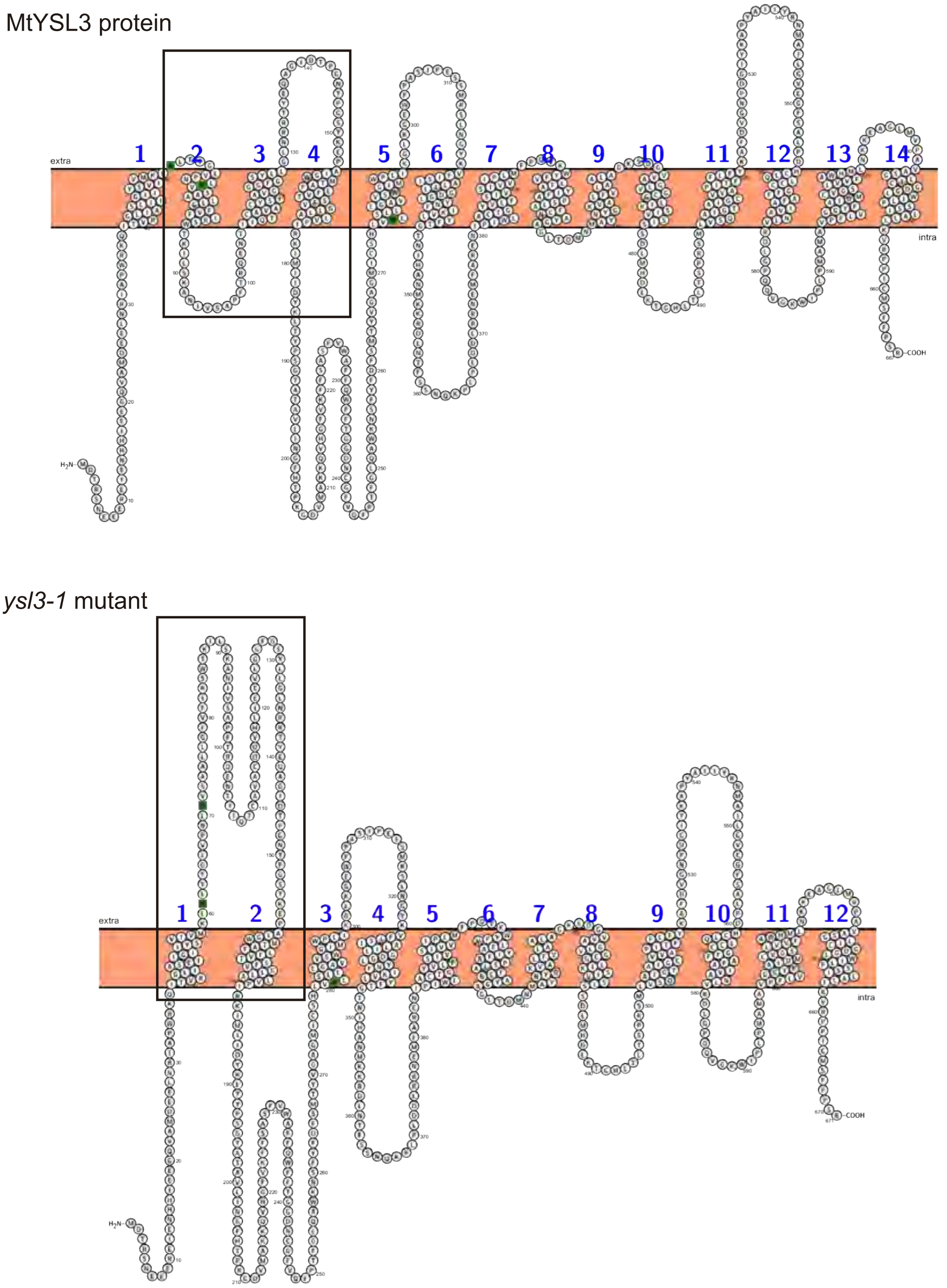


**Fig. S3. Effect of *ysl3-1 Tnt1* insertion in MtYSL3 topology.** A) MtYSL3 protein sequence in wild type plants (WT) compared with MtYSL3 protein in *ysl3-1 -/-* mutant. B) Predicted transmembrane domains in the wild type MtYSL3 protein compared with MtYSL3 protein in y*sl3-1 -/-* mutants.

**FIGURE S4**

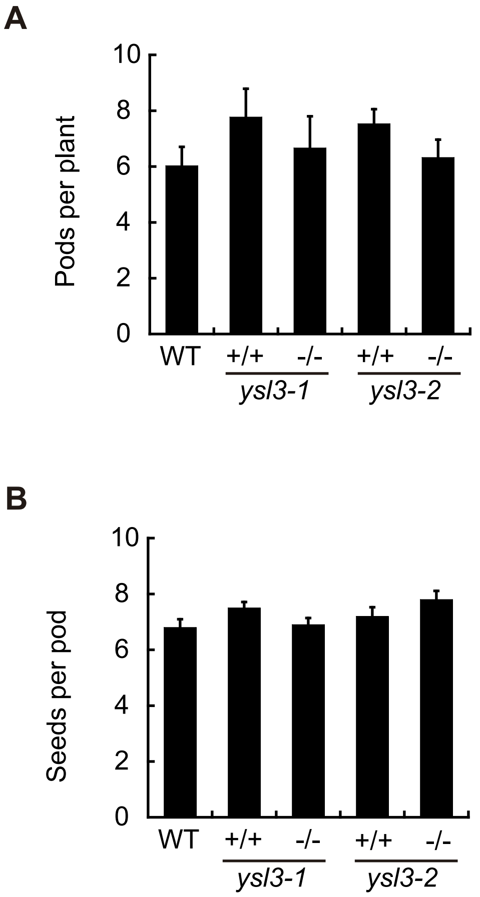


**Fig. S4. Effect of *MtYSL3* mutation in plant fertility under non-symbiotic conditions.** A) Average number of pods per plant. Data are the mean ± SE (n = 4 -10 plants). B) Average number of seeds per pod. Data are the mean ± SE (n = 15 pods).

**FIGURE S5**

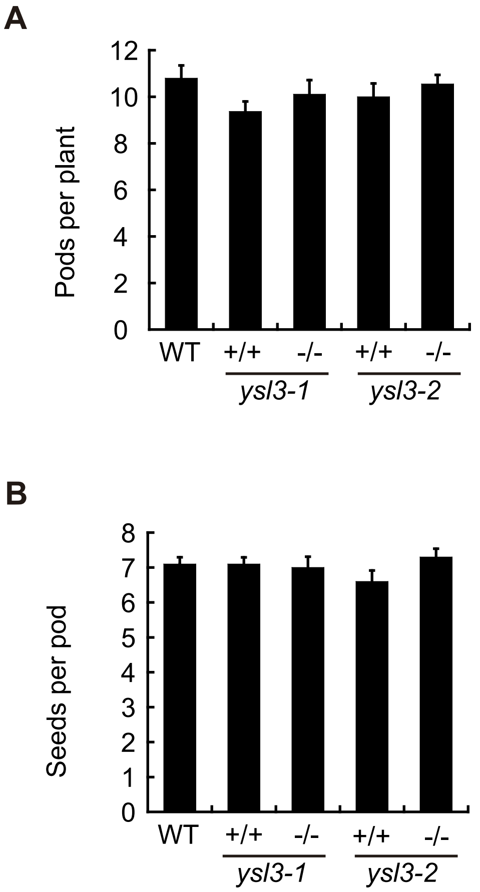


**Fig. S5. Effect of *MtYSL3* mutation in plant fertility under symbiotic conditions.** A) Average number of pods per plant. Data are the mean ± SE (n = 6-12 plants). B) Average number of seeds per pod. Data are the mean ± SE (n = 18-38 pods).

**FIGURE S6**

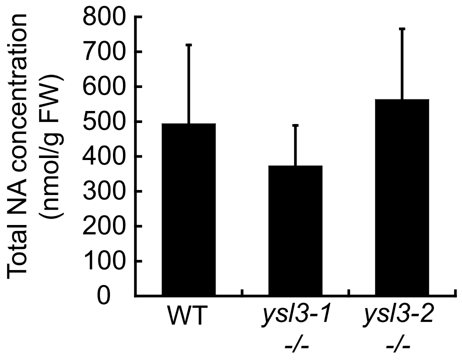


**Fig. S6. Nicotianamine content in 28 dpi wild type, *ysl3-1*, and *ysl3-2* nodules.** Data are the mean ± SE from five sets of nodules pooled from 8-10 plants.

Table S1. Primers used in this study

| Name | Sequence | Use |
|---|---|---|
| MtUb v4qF | ATTCTTCACATGCGGCGATTTAC | Quantitative expression of *MtUbiquitin carboxyl-terminal hydrolase* |
| MtUb v4qR | TTTCTCATTTGCTTTTGGTGTTGG | Quantitative expression of *MtUbiquitin carboxyl-terminal Hydrolase* |
| YSL2+2346qF_v2 | GGTTAGCATGTGTAGGGTATGTGGGAC | Quantitative expression of *MtYSL2* |
| YSL2+2467qR | CGGCACCATAAGCATTGCAGAAG | Quantitative expression of *MtYSL2* |
| YSL4qF_v2 | CAGTGCTCACCATAATATCCATCATTGTG | Quantitative expression of *MtYSL4* |
| YSL4qR_v2 | CGGCACCATAAGCATTGCAAAAA | Quantitative expression of *MtYSL4* |
| YSL3qF_v2 | GTGTGGGCAGTTTGGTTGTGTTTGC | Quantitative expression of *MtYSL3* |
| YSL3qR | GGGAAGAAGCTCATGCATATTGGGG | Quantitative expression of *MtYSL3* |
| YSL1qF | CTCCATCAAGTCCACTAAACATCAAC | Quantitative expression of *MtYSL1* |
| YSL1qR | ATGTGTTCTGTCCAATGTTCATCGG | Quantitative expression of *MtYSL1* |
| 5MtYSL3-2038GW | GGGGACAAGTTTGTACAAAAAAGCAGGCTCCGCTAATAAAATCAAATCAGT | *MtYSL3* promoter cloning into pGWB3 or pGWB13 Gateway vectors |
| 3MtYSL3-20GW | GGGGACCACTTTGTACAAGAAAGCTGGGTATGGATTCACAGATTCCACCAC | *MtYSL3* promoter cloning into pGWB3 |
| 3MtYSL3+3259GW | GGGGACCACTTTGTACAAGAAAGCTGGGTTCCTAGATGGGAAGAAGCTCAT | *MtYSL3* promoter and genomic region cloning into pGWB13 |
